# Cross-family Signaling: PDGF Mediates VEGFR Activation and Endothelial Function

**DOI:** 10.1101/2025.02.27.640684

**Authors:** Xinming Liu, Yingye Fang, Shreya Karra, Silvia Gonzalez-Nieves, Simon Guignard, Vincenza Cifarelli, P.I. Imoukhuede

## Abstract

Cross-family signaling between platelet-derived growth factors (PDGFs) and vascular endothelial growth factor receptors (VEGFRs) is a newly discovered yet unexplored mechanism of angiogenesis regulation. This study elucidates the role of PDGFs in endothelial cell (EC) signaling and functions, focusing on VEGFR1 and VEGFR2 activation. Using human dermal microvascular ECs (HDMECs) with double knockout of PDGFRα/β and human brain microvascular ECs (HBMECs), we show three key findings: (1) PDGF-AA and -BB induced VEGFR1 phosphorylation, peaking at 2-fold increases at low concentrations (0.5 ng/mL), while PDGF-AB stimulated a 2-fold rise in VEGFR2 phosphorylation. (2) Downstream effectors PLCγ1, Akt, and FAK were activated by all three PDGFs at levels comparable to VEGF-A. (3) PDGF-BB significantly enhanced EC proliferation (up to 240%) and migration (up to 170%), with lower PDGF concentrations (0.5–5 ng/mL) eliciting stronger effects than higher concentrations (50–100 ng/mL). Overall, PDGF subtypes differentially induce VEGFR phosphorylation, downstream effector activation, and angiogenic hallmarks such as proliferation and migration, revealing novel mechanisms for regulating endothelial function.

## Introduction

Angiogenesis, the formation of new blood vessels from existing vascular systems, is central to several physiological conditions, supporting tissue growth during development and wound healing, as well as in pathological conditions such as cancer [1]. The process of angiogenesis is primarily regulated by vascular endothelial growth factors (VEGFs) and platelet-derived growth factors (PDGFs), which bind to their respective receptors, VEGFRs and PDGFRs [2, 3]. This binding, described by the uni-family model, triggers downstream angiogenic signaling [4]. Dysregulated angiogenesis is associated with over 70 vascular diseases and cancers [5, 6]. Current anti-cancer therapies targeting VEGFs to inhibit excessive angiogenesis have demonstrated limited efficacy [7]. The anti-VEGF-A antibody has led to transient improvements, such as tumor stasis and shrinkage. However, after a period of weeks to months, the tumors typically resume growth and progress [8]. Surprisingly, tumors resistant to anti-VEGF treatment showed PDGF overexpression [7–9]. This finding suggests the involvement of other molecules in the pathological process beyond the commonly recognized uni-family VEGF-VEGFR signaling axis.

A novel paradigm of cross-family signaling has been proposed and identified between VEGF and PDGF families [4]. The two families share similarities structurally and functionally for both ligands (VEGFs and PDGFs) and receptors (VEGFRs and PDGFRs) [10–13]. The ligands bind to and activate their respective receptors in a similar way, followed by dimerization and tyrosine phosphorylation [2, 14, 15]. Studies in literature have established evidence of cross-family signaling. The bindings of VEGF-to-PDGFR and PDGF-to-VEGFR were both discovered with high affinities [4], indicating a complex interplay between these growth factor families. VEGF-A has shown the ability to activate PDGFRs in a dose-dependent manner, independently of VEGFRs [16]. Investigations in human dermal fibroblasts lacking VEGFRs have revealed that VEGF-A can modulate surface PDGFR levels and inhibit PDGF ligand interactions [16, 17]. Furthermore, PDGF-CC, a homodimer structurally similar to VEGF-A [18], has displayed the capacity to stimulate EC migration at a level comparable to VEGF-A [19]. Treatment of ECs with PDGFs has been associated with reduced VEGFR2 expression [20]. Computational models have predicted that PDGFs contribute significantly to VEGFR2 ligation in both healthy conditions and breast cancer [4]. Taken together, these data underscore the potential impact of VEGF-PDGF cross-family signaling in angiogenesis regulation. However, the resulting functional effects of PDGF-to-VEGFR binding have not been fully elucidated.

In this study, we aimed to investigate the role of PDGFs in regulating endothelial signaling and function via VEGFR1 and VEGFR2 by employing several *in vitro* strategies. To exclude the effects of PDGF signaling through PDGFRs, we engineered endothelial cells (ECs) to lack PDGFR-α and -β expression on the plasma membrane. As comparative control, we also used human dermal fibroblasts (HDFs) that lack VEGFRs but express PDGFRs to elucidate the functional effects of PDGF signaling through PDGFRs in HDFs and VEGFRs in ECs. To exclude the possible VEGF upregulation simulated by PDGF treatment, we validated the VEGF-A protein concentrations in the cell culture supernatant followed by PDGF stimulations compared with untreated conditions. Our comprehensive investigation assessed the activation of VEGFRs and their downstream effectors, i.e., PLCγ1, Akt, and FAK, and examined the resulting functional roles in angiogenesis, such as cell proliferation and migration. Through our research, we aim to shed light on alternative mechanisms that could potentially address drug resistance issues observed in anti-VEGF therapies, offering new insights into overcoming vascular dysregulation.

## Material and Methods

### Cell culture

Primary human brain microvascular endothelial cells (HBMECs) (Cell Systems, ACBRI 376) were cultured in Complete Classic Medium (Cell Systems, 4Z0-500) supplemented with CultureBoost and Bac-Off Antibiotic (Cell Systems, 4Z0-644). Primary human dermal microvascular endothelial cells (HDMECs) (ATCC, CRL-4060) double knock out (DKO) for *PDGFRA* and *PDGFRB* (to eliminate PDGFRα and β expression, respectively) were obtained via CRISPR/Cas9 at the Genome Engineering & Stem Cell Center at Washington University in St. Louis, MO. The efficiency of DKO was validated by next-generation sequencing analysis [21]. HDMECs (*PDGFRA*^*−/−*^ and *PDGFRB*^*−/−*^) were cultured in Vascular Cell Basal Medium (ATCC, PCS100030) supplemented with microvascular endothelial cell growth kit -VEGF (i.e., with VEGF; ATCC, PCS110041) and penicillin streptomycin (10,000 U/mL) (Thermo Fisher Scientific, 15-140-122). Human dermal fibroblasts (HDFs) were purchased from ATCC (PCS-201-010) and cultured in DMEM (Thermo Fisher Scientific, 11995065) with 5% FBS (Sigma-Aldrich, 12306C) and Penicillin Streptomycin (10,000 U/mL) (Thermo Fisher Scientific, 15-140-122). Cells were used between passages 3–6 and maintained in a humidified incubator at 37 °C and with 5% CO_2_. Low-serum medium used in this study for cell starvation was made from EBM-2 (Lonza, CC-3156) supplemented with 1% FBS, Bac-Off Antibiotic (Cell Systems, 4Z0-644), and necessary components minus growth factors (hydrocortisone, ascorbic acid, GA-1000, heparin).

### Growth factor application

The recombinant hPDGF-AA (R&D Systems, 221-AA), hPDGF-AB (R&D Systems, 222-AB), hPDGF-BB (R&D Systems, 220-BB), and hVEGF-A (R&D Systems, 293-VE/CF) were reconstituted with sterile 4 mM HCL (R&D Systems, RB03) at 100 μg/mL, verified by NanoDrop Spectrophotometer (ThermoFisher, ND-ONE-W). They were aliquoted and stored at -20 °C.

#### Proliferation assay

Cells were seeded in a 96-well plate and grown to approximately 50% confluence and serum-starved overnight using low-serum medium. Cells were stimulated with specific ligands, PDGF-AA, -AB, -BB, or VEGF-A (R&D Systems, 221-AA-050, 222-AB-050, 220-BB-050, 293-VE-050/CF; same materials were used throughout the study) at specific concentrations for 24 hours. XTT (2,3-Bis-(2-Methoxy-4-Nitro-5-Sulfophenyl)-2H-Tetrazolium-5-Carboxanilide) (Invitrogen, X6493) was dissolved in EBM-2 (Lonza, CC-3156) at 1 mg/mL and activated by PMS (N-methyl dibenzopyrazine methyl sulfate) (0.0075 mg/mL). The reagent was added to each well and mixed with pipetting. The plates were incubated at 37 °C for 3 hours in the dark. The absorbance was read at 450 and 650 nm. The final OD value is [Abs_450nm_(Test) – Abs_450nm_(Blank)] – Abs_660nm_(Test) and normalized to the untreated control.

#### Migration assay

Cells were cultured in T-75 flasks until they reached approximately 90% confluence. Subsequently, the cells were serum-starved overnight using low-serum medium. Transwell inserts (Falcon, 353097) were coated with 2.45 μg/mL Collagen type I (Corning, 354236) in PBS and incubated overnight at 4°C. After starvation, the cells were detached and seeded in the top chamber of the transwell inserts at a density of 1 ×10^5^ cells/well. The bottom chamber contained growth factors at the specified concentrations in low-serum medium. The cultures were then incubated for 24 hours at 37 °C with 5% CO_2_. Following incubation, the unmigrated cells on the upper surface of the chamber were gently removed using a cotton swab. The inserts were subsequently washed with PBS. The cells that had migrated through the membrane to the lower side of the insert were fixed with 4% paraformaldehyde, permeabilized with 0.1% Tween-20, and stained with a 0.01% Hoechst 3342 solution (Invitrogen, H3570). After a final wash with PBS, the inserts were imaged under the DAPI channel (EVOS M5000). Cell migration was quantified using ImageJ (1.52a) as the number of migrated cells to the downside of the inserts.

#### Enzyme-Linked Immunosorbent Assay (ELISA) / Immunoblotting for phosphorylation detection

Cells (∽90% confluent) were serum starved for 6 hours using low-serum medium. The indicated growth factors were added into the medium and incubated for 30 minutes for detecting phospho-VEGFR1 in ECs and phospho-PDGFRβ in HDFs or 10 minutes for detecting phospho-VEGFR2 in ECs. Treatment medium was aspirated to terminate the stimulation, and the plates were washed with ice-cold phosphate-buffered saline (PBS). For ELISA, cell lysates were collected with lysis buffer (R&D Systems, DYC002) with protease and phosphatase inhibitor cocktail (Sigma-Aldrich, P8340). After being briefly vortexed and centrifuged at 12,000×*g* for 10 min at 4 °C, the cell lysates were used to assess the phosphorylation levels of VEGFR1, VEGFR2, or PDGFRβ, respectively using Human Phospho-VEGFR1/Flt-1 DuoSet IC ELISA kit (R&D Systems, DYC4170), Human Phospho-VEGFR2/KDR DuoSet IC ELISA kit (R&D Systems, DYC1766), or Human Phospho-PDGFRβ DuoSet IC ELISA kit (R&D Systems, DYC1767) per manufacturer’s instruction. The total protein concentrations were quantified through bicinchoninic acid assay (BCA) (Pierce Biotech, 23227). The levels of phosphorylation were normalized to untreated control samples.

For immunoblotting, the cells were scraped in ice-cold 1× RIPA lysis buffer (EMD Millipore, 20-188) supplemented with protease and phosphatase inhibitor cocktails (Sigma-Aldrich, 11836170001 and 4906837001). The cell lysates were cleared by centrifuge at 12,000×*g* for 10 min at 4 °C, mixed with 6 × SDS buffer (Boston BioProducts, BP-111R), denatured at 95 °C for 10 min, resolved on 4–15% SDS-PAGE gels (Bio-Rad, 4561084), and transferred to nitrocellulose membranes (Bio-Rad, 1620215). The membranes were blocked with 5% non-fat dry milk dissolved in 1× Tris Buffered Saline (TBS) for 1 hour at room temperature before adding primary antibodies (Supplementary Table 1) overnight at 4 °C. Secondary antibodies were added and incubated for 20 min at room temperature. Protein signals were detected using the LI-COR Odyssey CLx Imager (LI-COR Biosciences) and quantified using ImageJ Software (2.14.0/1.54f) or Image Studio 6.0 (LI-COR Biosciences). The levels of phosphorylation were normalized to total VEGFR2 proteins and to untreated control samples.

#### Enzyme-Linked Immunosorbent Assay (ELISA) for VEGF-A secretion detection

Cells (∽90% confluent) were serum starved for 6 hours using low-serum medium. The indicated growth factors were added into the medium and incubated for 10 min, 30 min, or 24 hours. The cell culture supernatant was collected and centrifuged at 1000×*g* for 5 min. The concentration of VEGF-A in the cell culture supernatant was detected with Human VEGF ELISA Kit - Quantikine (R&D Systems, SVE00) following the instructions. The results were normalized to untreated control samples and statistically analyzed.

#### Quantitative flow cytometry

ECs were seeded in T-75 flasks and grown to approximately 80% confluence. The cells were washed with Ca^2+^/Mg^2+^-free PBS, harvested in Corning CellStripper solution, a nonenzymatic cell-dissociation solution, and incubated for 15 min at 37 °C and 5% CO_2_. The disassociation was stopped by adding Ca^2+^/Mg^2+^-free PBS, and the cell suspension was centrifuged twice at 400×*g* for 5 min at 4 °C. The cells were resuspended at 2–4 ×10^6^ cells/mL in ice-cold stain buffer (PBS, bovine serum albumin, sodium azide). A single-cell suspension was added for plasma membrane staining (25 μL) or for whole-cell staining (50 μL) to 96-well deep-well plates. Human VEGFR1 phycoerythrin (PE)-conjugated antibody (R&D Systems, FAB321P), VEGFR2 PE-conjugated antibody (BioLegend, 359904), VEGFR3 PE-conjugated antibody (R&D Systems, FAB3492P), PDGFRα PE-conjugated antibody (R&D Systems, FAB1264P), or PDGFRβ PE-conjugated antibody (R&D Systems, FAB1263P) was added to each sample well except the fluorescence-minus-one (FMO) control. The saturating concentrations of PE-conjugated antibodies to detect receptors on the plasma membrane were previously determined at 14 μg/mL for VEGFR1, 2, and 3; and 9.4 μg/mL for PDGFRα and PDGFRβ [17, 22]. For whole-cell receptor detection, the saturating concentration of VEGFR1 and VEGFR2 antibodies was determined at the beginning of the study to be 20 ug/mL for both, and PE-VEGFR3 antibody saturated at 30 μg/mL [21]. For PDGFR antibodies the same concentration was used as that of VEGFR antibodies. Samples were incubated in dark for 40 min at 4 °C and washed twice with stain buffer by centrifugation at 400×*g* for 5 min at 4 °C. Samples for plasma membrane receptor measurement were washed and resuspended in 100 μL stain buffer before analysis.

Samples for whole-cell receptor measurement were fixed using 2% paraformaldehyde for 20 min in the dark, washed, and permeabilized (0.5% Tween-20) for 20 min in the dark. Samples were washed twice with stain buffer containing 0.1% Tween-20. PE-conjugated antibodies were added to the samples for staining for 40 min in the dark, followed by three washes with stain buffer containing 0.1% Tween-20. Washed samples for whole-cell receptor measurement were resuspended in 50 μL stain buffer. The readouts were acquired by a Cytoflex S flow cytometer (Beckman Coulter, IN) and CytExpert software (Beckman Coulter, IN). Kaluza software (Beckman Coulter, IN) was used for data analysis. Quantibrite PE beads (BD Biosciences, 340495) were collected and analyzed under the same compensation and voltage settings as cell fluorescence data. Quantibrite PE beads are a mixture of beads with four different concentrations of conjugated PE molecules: low (474 PE molecules/bead), medium-low (5359 PE molecules/bead), medium-high (23,843 PE molecules/bead), and high (62,336 PE molecules/bead). A calibration curve that translated PE geometric mean to the number of bound molecules was determined using linear regression, *y* = *mx* + *c*, where y is log_10_ fluorescence and x is log_10_ PE molecules per bead. Receptor levels were calculated as described previously [22–25].

#### Statistics

All data were acquired in at least three independent experiments. Ensemble averaged data are represented by mean ± standard error of the mean (SEM). One-way analysis of variance (ANOVA), multiple comparison post-hoc Tukey’s test, and power analysis of pair-comparison were performed.

## Results

### Selection of VEGFR-expressing cell lines for investigating PDGF-induced signaling and functions

We assessed VEGFR levels on the plasma membrane and across the whole cells on five different human EC lines. HBMECs were selected based on their highest plasma membrane VEGFR2 expression and comparable VEGFR1 levels among five EC lines, making them optimal for investigating PDGF-induced VEGFR signaling (Figure 1A). Further, to eliminate potential PDGF signaling through PDGFRs, we engineered a *PDGFRA* and *PDGFRB* double knockout (PDGFR-DKO) human dermal microvascular endothelial cell (HDMEC) line using CRISPR/Cas9. HDMECs (*PDGFRA*^*−/−*^ and *PDGFRB*^*−/−*^) expressed <100 PDGFRα/cell and <100 PDGFRβ/cell on the plasma membrane (Figure 1C), which likely resulted from nonspecific binding as described previously [24, 25]. Similarly, HBMECs exhibited negligible amounts of plasma membrane PDGFRs (<300 PDGFRs/cell) when compared to HDMECs (*PDGFRA*^*−/−*^ and *PDGFRB*^*−/−*^) (p < 0.05). The whole-cell PDGFR expressions were about 10 times higher than the expression levels on the plasma membrane, likely because of increased nonspecific binding during whole-cell staining (Figure 1D). As a control, we performed the same PDGF treatments on human dermal fibroblast (HDF) cells, which express PDGFRs but lack VEGFRs on their plasma membranes [26], to confirm the PDGF protein activity.

**Figure 1.**
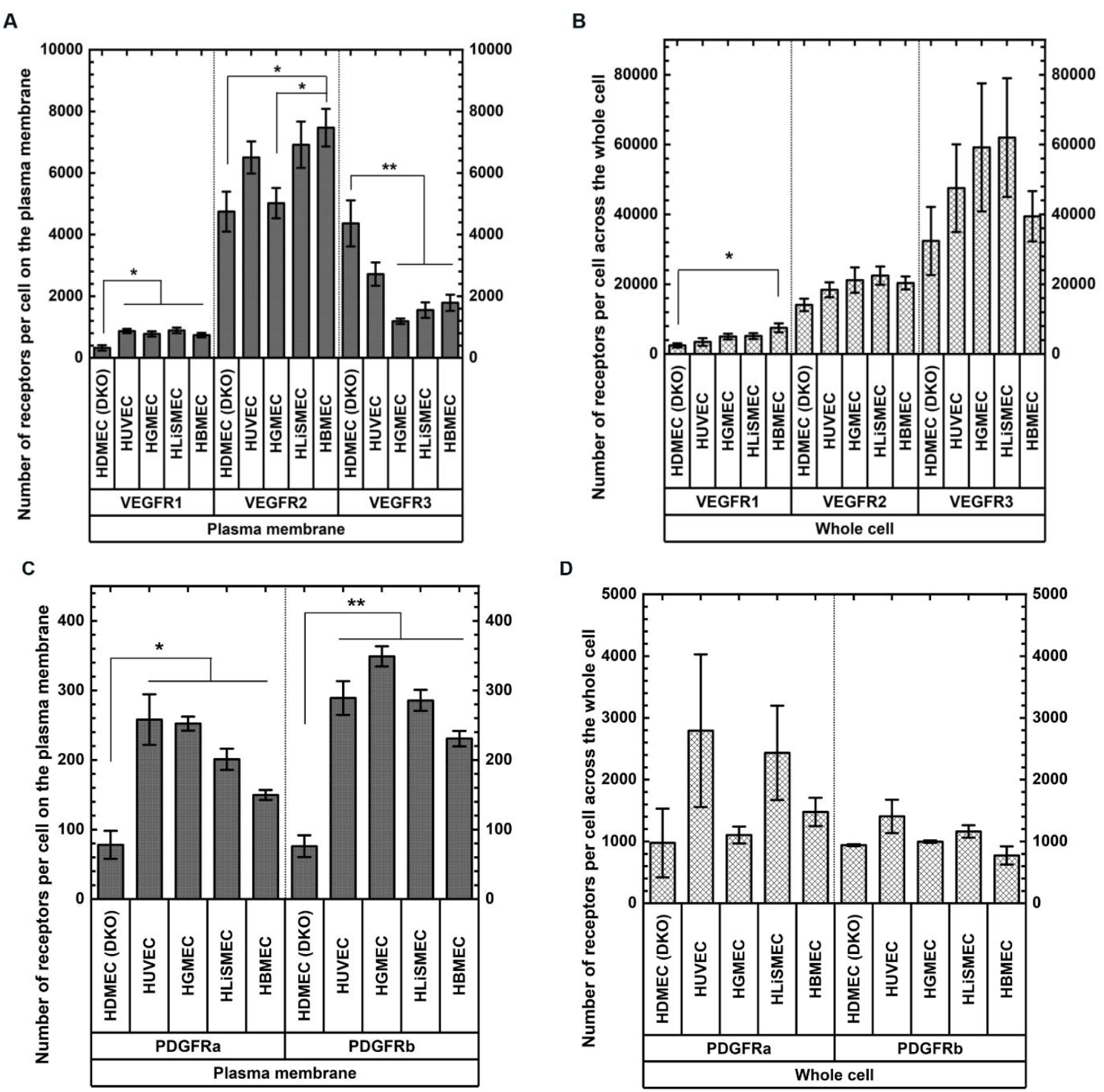
Levels of VEGFRs and PDGFRs on the plasma membrane and in the whole cells of human ECs from different origins. ECs from different organs presented different VEGFR levels **(A)** on the plasma membrane and **(B)** in the whole cells assessed by quantitative flow cytometry. Expression levels of PDGFRα or PDGFRβ either **(C)** on the plasma membrane or **(D)** in the whole cells are negligible across five different human EC cell lines. Error bars represent the standard error of mean (SEM). *p<0.05, **p<0.01, ***p<0.001 by one-way ANOVA and multi-comparison post-hoc Tukey’s test.

### PDGF-AA and -BB induced VEGFR1 phosphorylation, whereas PDGF-AB phosphorylated VEGFR2

We initially assessed receptor activation by quantifying VEGFR1 and VEGFR2 phosphorylation through ELISA and immunoblotting following growth factor stimulation. We treated HBMECs and HDMECs (*PDGFRA*^*−/−*^ and *PDGFRB*^*−/−*^) with PDGF-AA, -BB, and -AB at increasing concentrations from 0 to 100 ng/mL. HBMECs showed an overall descending trend as PDGF-AA and -BB treatment concentrations increased (Figure 2A, 2C). We observed that 0.5 ng/mL PDGF-AA and PDGF-BB treatments significantly increased VEGFR1 phosphorylation and presented a peak by a 1.8 and 2.2-fold increase, respectively, compared with untreated HBMECs (p<0.001), which was similar to the changes of VEGF-A treatment (Figure 2A, 2C). PDGF-AB did not have an effect on VEGFR1 phosphorylation (Figure 2B). We did not observe any PDGF-induced VEGFR2 phosphorylation in HBMECs, regardless of low or high concentrations of PDGFs (Figure 2D, 2E).

**Figure 2.**
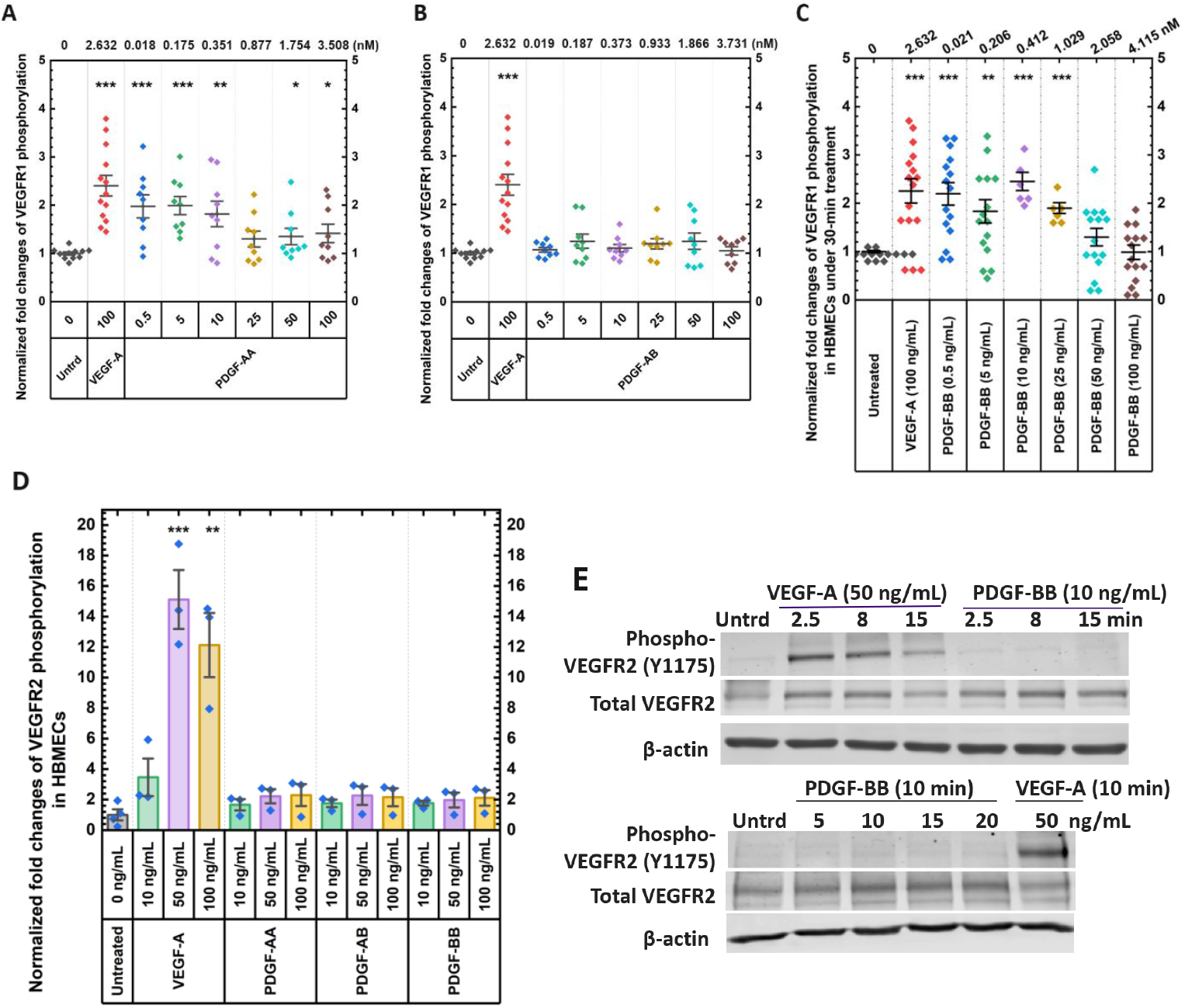
Low concentrations of PDGF-AA and -BB significantly phosphorylated VEGFR1 but not VEGFR2 in HBMECs. HBMECs were serum-starved for 6 hours and stimulated with growth factors at the indicated concentrations. Activation of VEGFR1 or VEGFR2 was measured either by ELISA detecting the pan-tyrosine phosphorylation of **(A–C)** VEGFR1 or **(D)** VEGFR2, or **(E)** by immunoblotting and probed with primary antibody targeting phospho-VEGFR2 at tyrosine site 1175. The concentrations of stimulating growth factors (ng/mL) are presented at the bottom of the plots, while the concentrations converted to the unit of nM are at the top of A–C. Three independent replicates were performed and analyzed. Error bars represent the standard error of mean (SEM). (*p<0.05, **p<0.01, ***p<0.001 as determined by one-way ANOVA and multi-comparison post-hoc Tukey’s test.)

In HDMECs (*PDGFRA*^*−/−*^ and *PDGFRB*^*−/−*^), VEGFR1 phosphorylation was not observed stimulated by PDGF-AA (Figure 3A), whereas PDGF-AB demonstrated a 1.8-fold increase at 10 ng/mL (p < 0.05) (Figure 3B). PDGF-BB at 25 ng/mL activated VEGFR1 to the greatest level to 180% (p < 0.001) (Figure 3C). Phosphorylation of VEGFR2 at tyrosine site 1175 was highly induced by 5 ng/mL PDGF-AB to 200% elevation after 15-min stimulation (Figure 3F), whereas the other two PDGFs showed no significant impacts (Figure 3E, 3G). VEGF-A induced the activation to a greater extent to 360%– 400% increase at a relatively high concentration of 50 ng/mL, aligning with the observations from ELISA conducted in HBMECs (Figure 3D, 2D).

**Figure 3.**
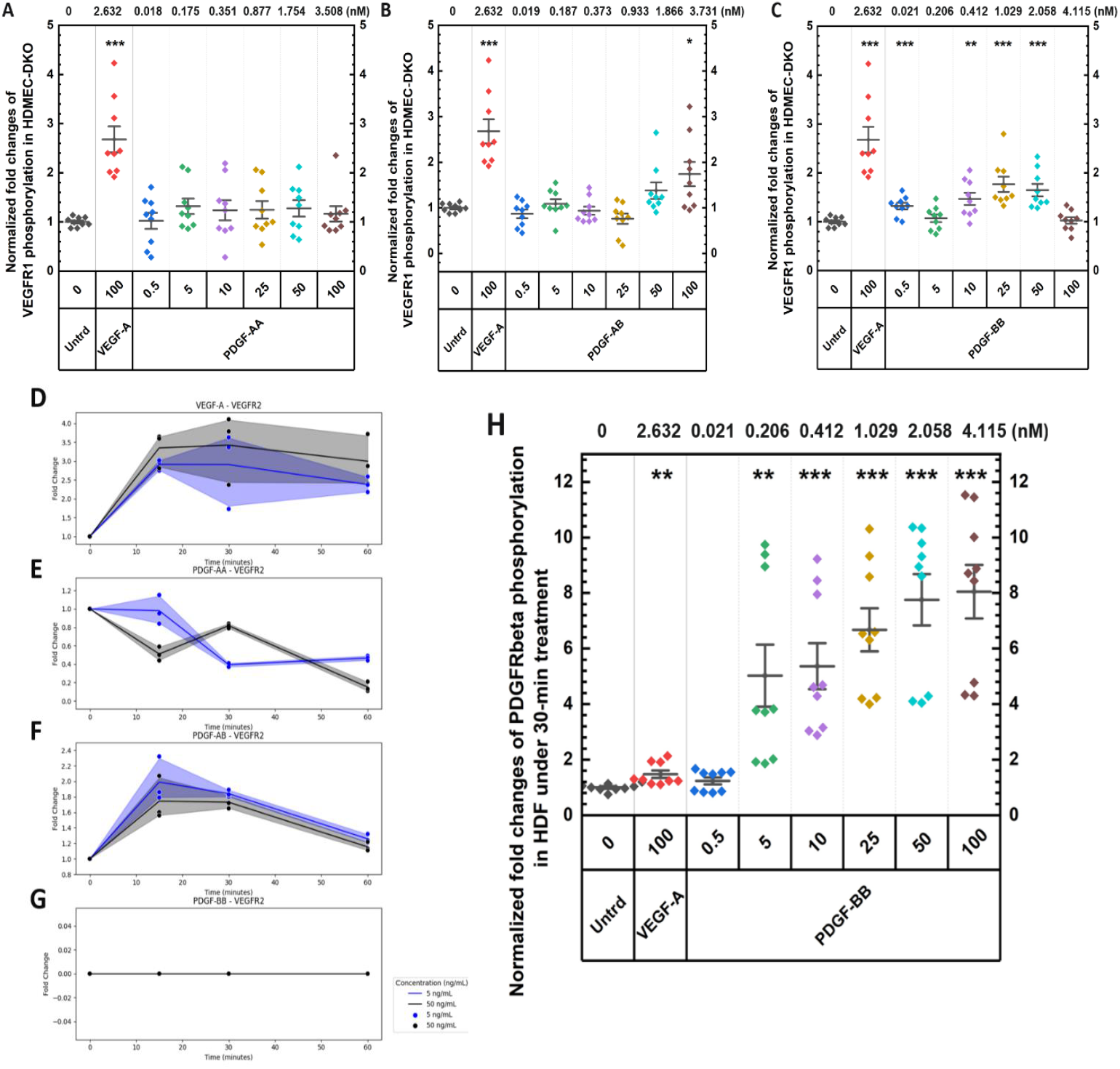
High concentrations of PDGF-BB induced VEGFR1 phosphorylation, whereas low concentrations of PDGF-AB activated VEGFR2 in HDMECs (*PDGFRA*^*−/−*^ and *PDGFRB*^*−/−*^). **(A– G)** HDMEC (*PDGFRA*^*−/−*^ and *PDGFRB*^*−/−*^) were serum-starved for 6 hours and stimulated with growth factors at the indicated concentrations. **(A–C)** The fold changes of VEGFR1 phosphorylation were assessed by ELISA. **(D–G)** The fold changes of VEGFR2 phosphorylation were assessed via immunoblotting. **(H)** The fold changes of PDGFRβ phosphorylation in HDFs induced by 30-min VEGF-A or PDGF-BB treatment were measured via ELISA. The concentrations of stimulating growth factors (ng/mL) are presented at the bottom of the plot, while the concentrations converted to the unit of nM are at the top of A–C and H. Three independent replicates were performed and analyzed. Error bars represent the standard error of mean (SEM). *p<0.05, **p<0.01, ***p<0.001 by one-way ANOVA and multi-comparison post-hoc Tukey’s test.

In HDFs, PDGFRβ phosphorylation was assessed to confirm PDGF-mediated activation of PDGFRβ. PDGF-BB induced robust PDGFRβ phosphorylation (Figure 3H). The significant effects were detectable at 5 ng/mL and peaked at 100 ng/mL, where the phosphorylation level was 8-fold higher compared with untreated cells. In contrast, treatment with 100 ng/mL VEGF or 0.5 ng/mL PDGF-BB resulted in low PDGFRβ phosphorylation, approximately 1.5-fold higher than control levels (Figure 3H).

### PDGFs activated adaptor proteins in HDMECs-DKO to a level comparable to that of VEGF-A

The effects of PDGF-AA, -AB, or -BB (5 and 50 ng/mL) on downstream protein activation (i.e., PLC γ1, Akt, and FAK) were assessed in HDMECs (*PDGFRA*^*−/−*^ and *PDGFRB*^*−/−*^) over time. Both PDGF-AA and -AB concentrations significantly phosphorylated PLCγ1 at tyrosine site 783 (p < 0.001) by similar fold changes to 200%–225% within 30-min treatment, which is comparable to the effects by 50 ng/mL VEGF-A (Figure 4A–4C). In contrast, 5 ng/mL of VEGF-A induced PLCγ1 phosphorylation to over 300% increase (p < 0.001) (Figure 4A). PDGF-BB, however, showed a subtle phosphorylation elevation to1.2-fold at 50 ng/mL after 60-min incubation (p < 0.05) (Figure 4D). The phosphorylation of Akt at serine site 473 showed the highest increase of all investigated growth factors to a 6–15-fold elevation (p < 0.001) (Figure 4E–4H).

**Figure 4.**
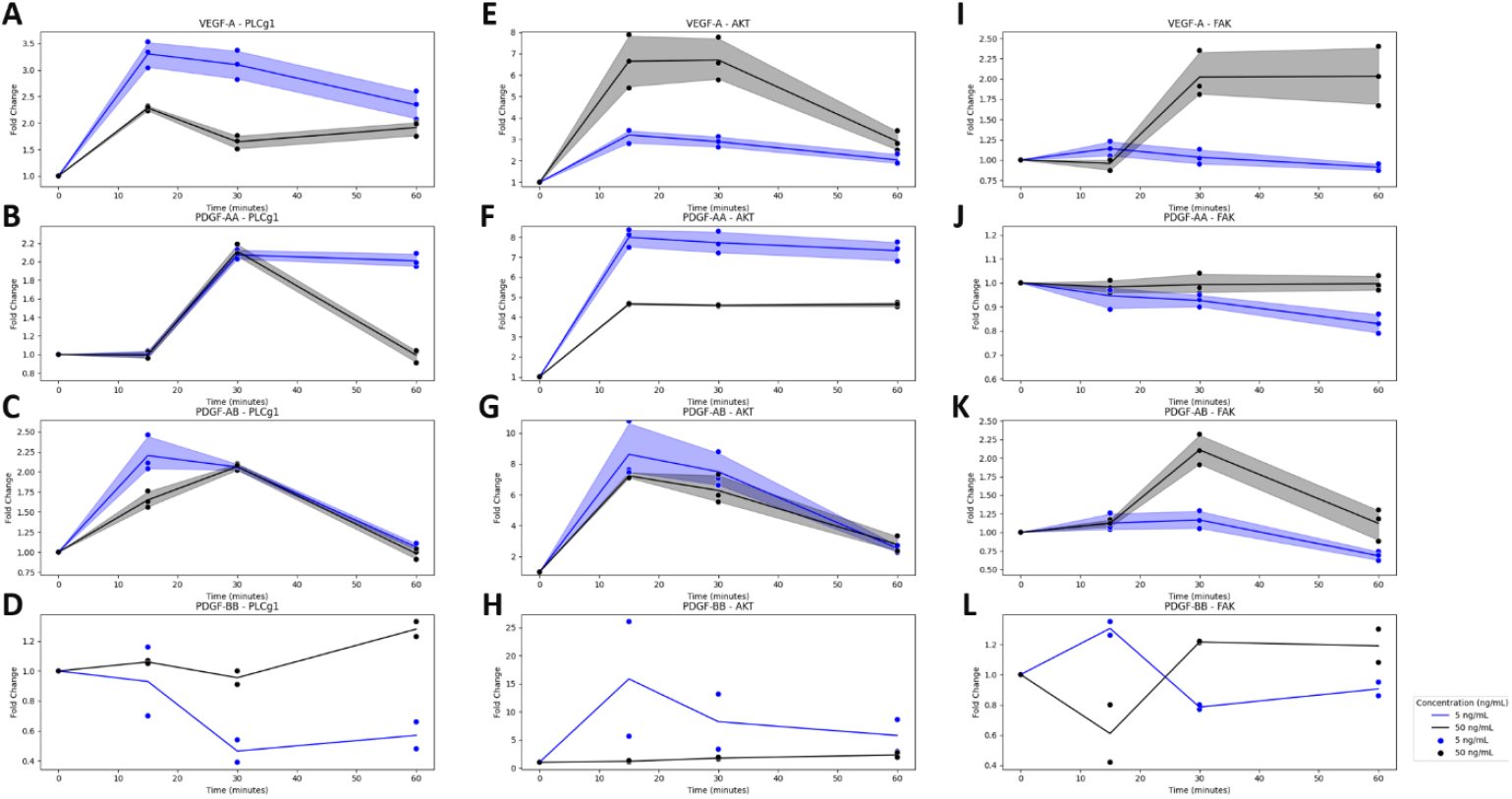
PDGFs activated adaptor proteins at levels comparable to those activated by VEGF-A. HDMECs (*PDGFRA*^*−/−*^ and *PDGFRB*^*−/−*^) were treated with PDGF-AA, -AB, or -BB at the concentrations of 5 and 50 ng/mL for 15 min. The effects of PDGF treatment on phosphorylating **(A-D)** PLCγ1 at tyrosine site 783, **(E-H)** Akt at serine site 473, and **(I-L)** FAK at tyrosine site 397 were assessed via immunoblotting.

Notably, PDGF-AA and -AB at 5 ng/mL showed Akt activation at a similar level to VEGF-A at 50 ng/mL, whereas PDGF-BB showed the highest level to 15-fold elevation at 5 ng/mL after 15-min treatment (Figure 4E–4H). Growth factors showed the lowest impacts on the activation of FAK at tyrosine site 397 with a 120–200% increase (Figure 4I–4L). A high concentration of 50 ng/mL PDGF-AB displayed a 2-fold increase in FAK phosphorylation (p < 0.001), comparable to that of 50 ng/mL VEGF-A after 30-min treatment (p < 0.05) (Figure 4I, 4K). PDGF-BB showed a minor increase at 1.2 times (p < 0.05), whereas PDGF-AA did not significantly phosphorylate FAK at the site (p = 0.86) (Figure 4J, 4L).

### PDGFs induced EC proliferation at levels comparable to VEGF-A

The activation of PLCγ1, Akt, and FAK has been found to modulate VEGF-induced EC proliferation, migration, and survival [27–30], which served as the foundation for exploring PDGF-induced EC proliferation. Serum-starved cells were treated with 0.5 ng/mL to 100 ng/mL VEGF-A or PDGFs. Notably, all PDGFs (PDGF-AA, -AB, and -BB) significantly stimulated EC proliferation. HBMECs treated with PDGF-AA or -AB exhibited up to a ∽150% increase in proliferation when the concentrations were between 0.5 ng/mL and 50 ng/mL (p < 0.01) (Figure 5A, 5B). PDGF-BB exhibited an inverse dose-response relationship: the low PDGF-BB concentration, 0.5 ng/mL, induced the highest HBMEC proliferation, resulting in a remarkable 240% increase compared with untreated cells (p < 0.001) (Figure 5C). As expected, VEGF-A treatment served as a positive control, demonstrating a substantial increase in cell growth, reaching up to 200% at concentrations ranging from 0.5 ng/mL to 100 ng/mL (Figure 5D). HDMEC (*PDGFRA*^*−/−*^ and *PDGFRB*^*−/−*^) showed significant increases in proliferation relative to untreated conditions across all investigated concentrations of PDGF or VEGF-A (p < 0.001). Notably, this increased proliferation was similar across all concentrations (Figure 6). In HDFs, PDGF-BB-mediated proliferation response in HDFs was the highest (p < 0.001) with up to a 280% increase in proliferation relative to the untreated control across the 5 ng/mL to 100 ng/mL concentration range (Figure 7A). Notably, VEGF-A mediated a significant proliferation response in HDFs (p < 0.001), relative to the untreated control, at concentrations between 10 ng/mL to 100 ng/mL, promoting proliferation by 250% (Figure 7B).

**Figure 5.**
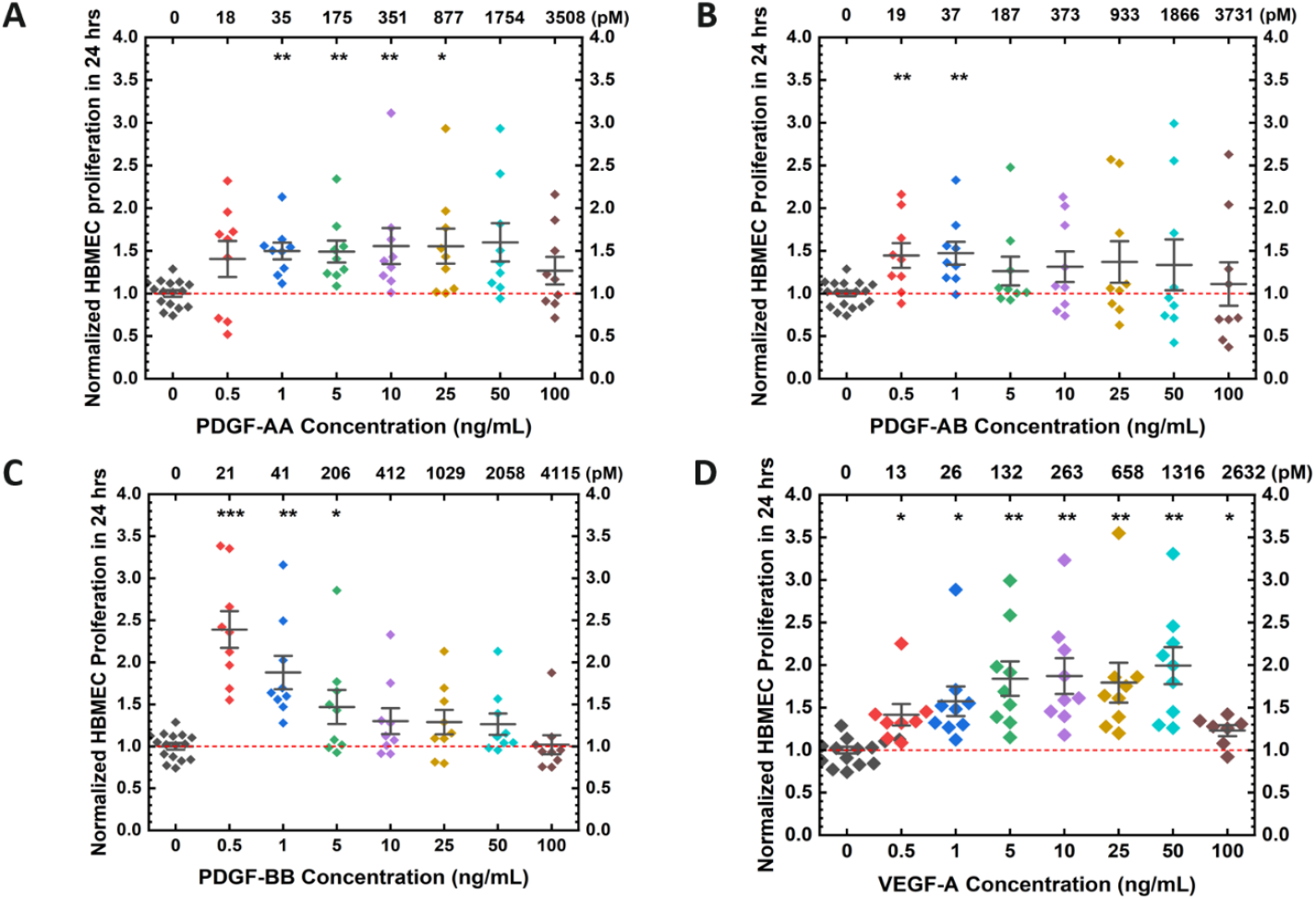
Low concentrations of PDGFs significantly induced HBMEC proliferation. HBMECs were treated with **(A)** PDGF-AA, **(B)** -AB, **(C)** -BB, or **(D)** VEGF-A with indicated concentration for 24 hours and subjected to proliferation assay. Cell viability was assessed using the XTT (2,3-Bis-(2-Methoxy-4-Nitro-5-Sulfophenyl)-2H-Tetrazolium-5-Carboxanilide) assay. OD value for each condition was normalized by untreated control. The concentrations of stimulating growth factors (ng/mL) are presented at the bottom of the plot, while the concentrations converted to the unit of pM are at the top of the plot. Three independent replicates were performed and analyzed. Error bars represent the standard error of mean (SEM). *p<0.05, **p<0.01, ***p<0.001 as determined by one-way ANOVA and multi-comparison post-hoc Tukey’s test.

**Figure 6.**
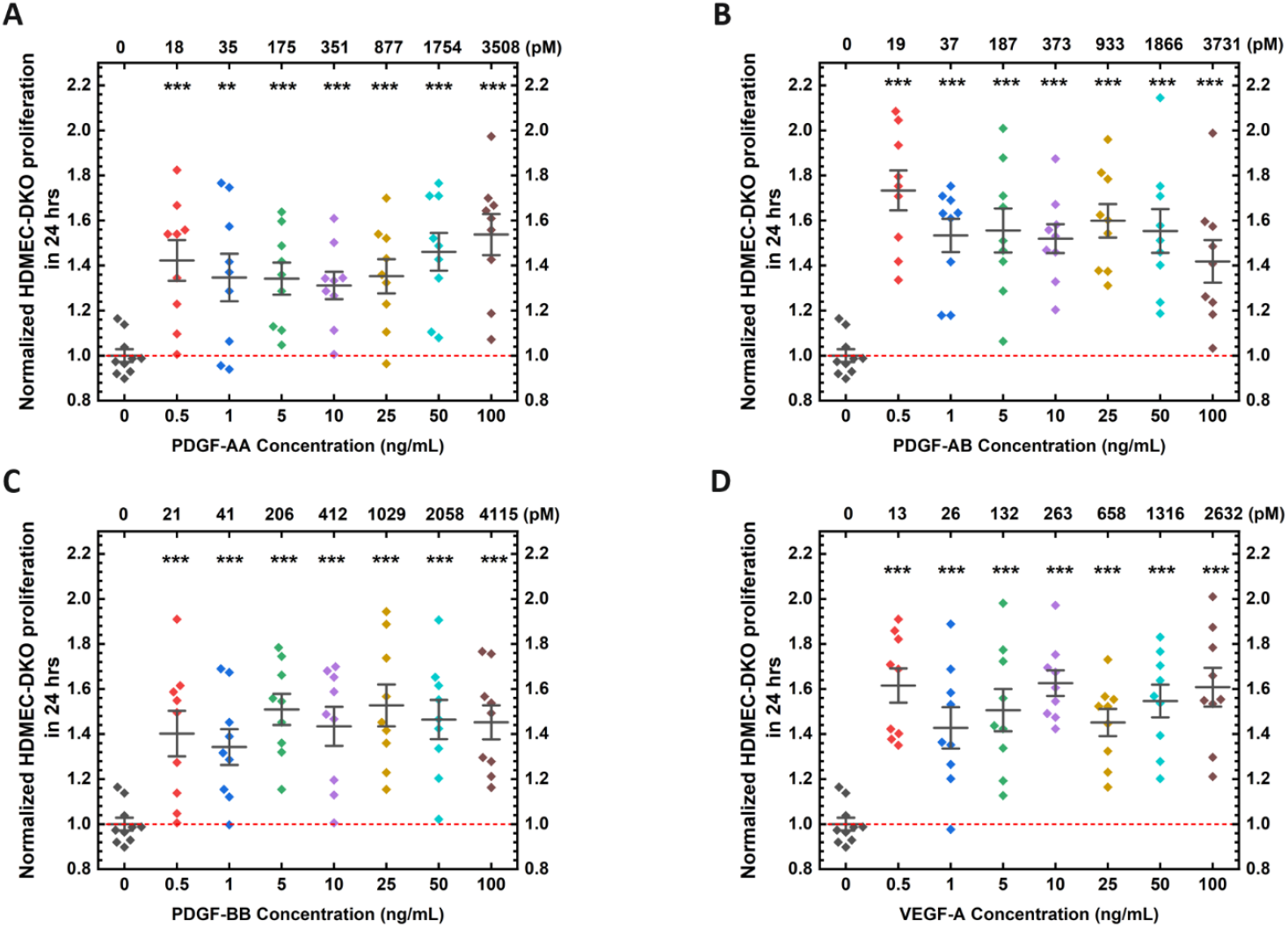
PDGFs induced proliferation of HDMECs (*PDGFRA*^*−/−*^ and *PDGFRB*^*−/−*^) at levels comparable to VEGF-A. HDMECs (*PDGFRA*^*−/−*^ and *PDGFRB*^*−/−*^) were treated with **(A)** PDGF-AA, **(B)** -AB, **(C)** -BB, or **(D)** VEGF-A with indicated concentration for 24 hours and subjected to XTT proliferation assay, as detailed in Material and Methods. The concentrations of stimulating growth factors (ng/mL) are presented at the bottom of the plot, while the concentrations converted to the unit of pM are at the top of the plot. Three independent replicates were performed and analyzed. Error bars represent the standard error of mean (SEM). *p<0.05, **p<0.01, ***p<0.001 as determined by one-way ANOVA and multi-comparison post-hoc Tukey’s test.

**Figure 7.**
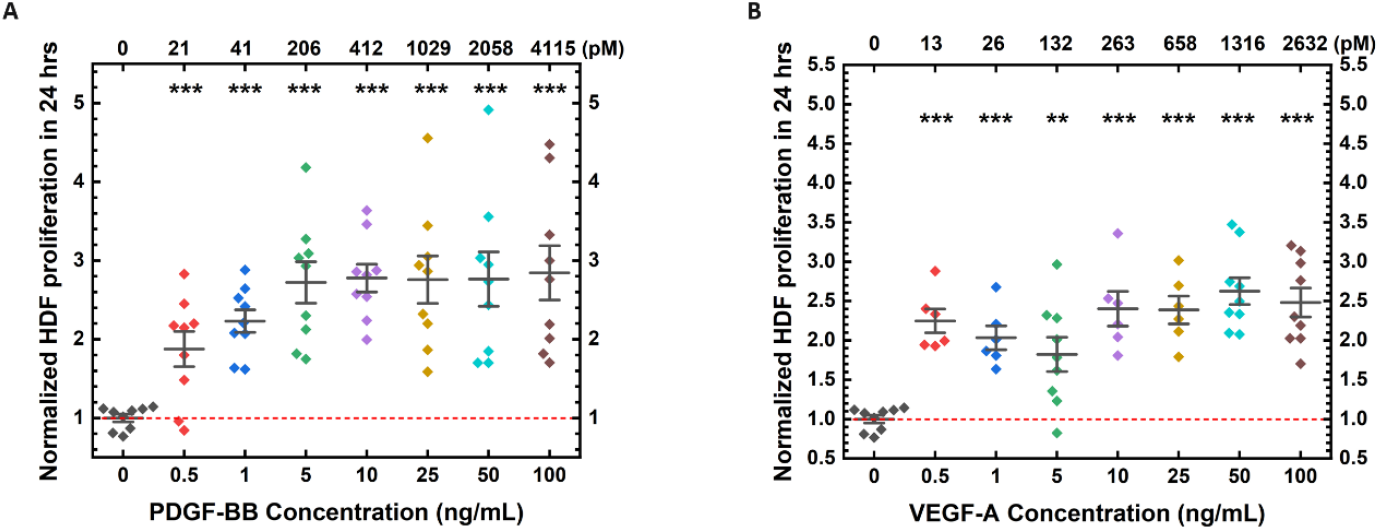
PDGF-BB induced a robust proliferation in human dermal fibroblast (HDF). HDFs were treated with **(A)** PDGF-BB, or **(B)** VEGF-A with indicated concentration for 24 hours and subjected to XTT proliferation assay, as detailed in Material and Methods. The concentrations of stimulating growth factors (ng/mL) are presented at the bottom of the plot, while the concentrations converted to the unit of pM are at the top of the plot. Three independent replicates were performed and analyzed. Error bars represent the standard error of mean (SEM). *p<0.05, **p<0.01, ***p<0.001 as determined by one-way ANOVA and multi-comparison post-hoc Tukey’s test.

### PDGF induced EC migration to 70% of the level stimulated by VEGF-A

To investigate the impact of PDGF on cell migration, we quantified the number of migrated cells through the transwell inserts stimulated by growth factors across the 0.5 ng/mL to 100 ng/mL concentration range for 24 hours. All investigated PDGF significantly triggered EC migration. In HBMECs, we observed that 25 ng/mL PDGF-AA or 10 ng/mL PDGF-AB induced the highest level of migration to 170% and 160% compared with untreated cells, respectively (p < 0.001) (Figure 8A, 8B). PDGF-BB increased HBMEC migration to 130% at 0.5 ng/mL (p < 0.001) (Figure 8C). In HDMEC (*PDGFRA*^*−/−*^ and *PDGFRB*^*−/−*^), low concentrations of PDGF-AA and -AB resulted in the most remarkable migration with a 130% enhancement relative to the untreated control at 5 ng/mL and 0.5 ng/mL, respectively (p < 0.01 and p < 0.001, respectively) (Figure 9A, 9B). PDGF-BB exhibited less migration with increasing concentrations in both EC types, with the lowest concentration, 0.5 ng/mL PDGF-BB resulting in a significant, 1.3-fold increase in migration relative to the untreated control in HDMECs (*PDGFRA*^*−/−*^ and *PDGFRB*^*−/−*^) (p < 0.001) (Figure 9C). Conversely, increasing PDGF-BB concentrations correlated with increasing migration of HDFs, starting from 5 ng/mL and peaking with a 2.8-fold increase in migration at 100 ng/mL (p < 0.001) (Figure 9D). VEGF-A served as a positive control with a substantial increase of up to 230% in both ECs (p < 0.001) but failed to show significant changes in HDFs (p = 0.09).

**Figure 8.**
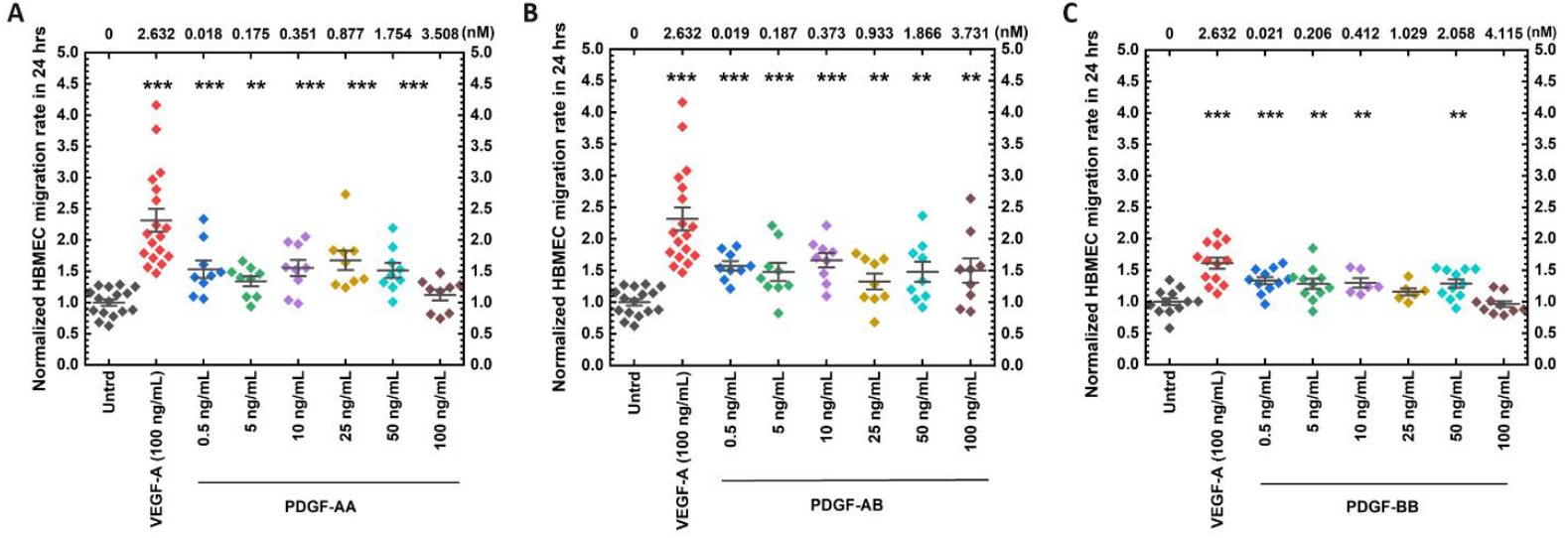
PDGFs induced migration of HBMEC lacking PDGFRs. HBMECs were seeded in the top chamber of transwell inserts and stimulated with **(A)** PDGF-AA, **(B)** -AB, **(C)** -BB, or VEGF-A at the indicated concentrations in low-serum medium for 24 hours. The cells that migrated through the membrane to the downside of the insert were fixed, permeabilized, stained, imaged, and counted on the following day. The concentrations of stimulating growth factors (ng/mL) are presented at the bottom of the plot, while the concentrations converted to the unit of nM are at the top of the plot. Three independent replicates were performed and analyzed. Error bars represent the standard error of mean (SEM). *p<0.05, **p<0.01, ***p<0.001 as determined by one-way ANOVA and multi-comparison post-hoc Tukey’s test.

**Figure 9.**
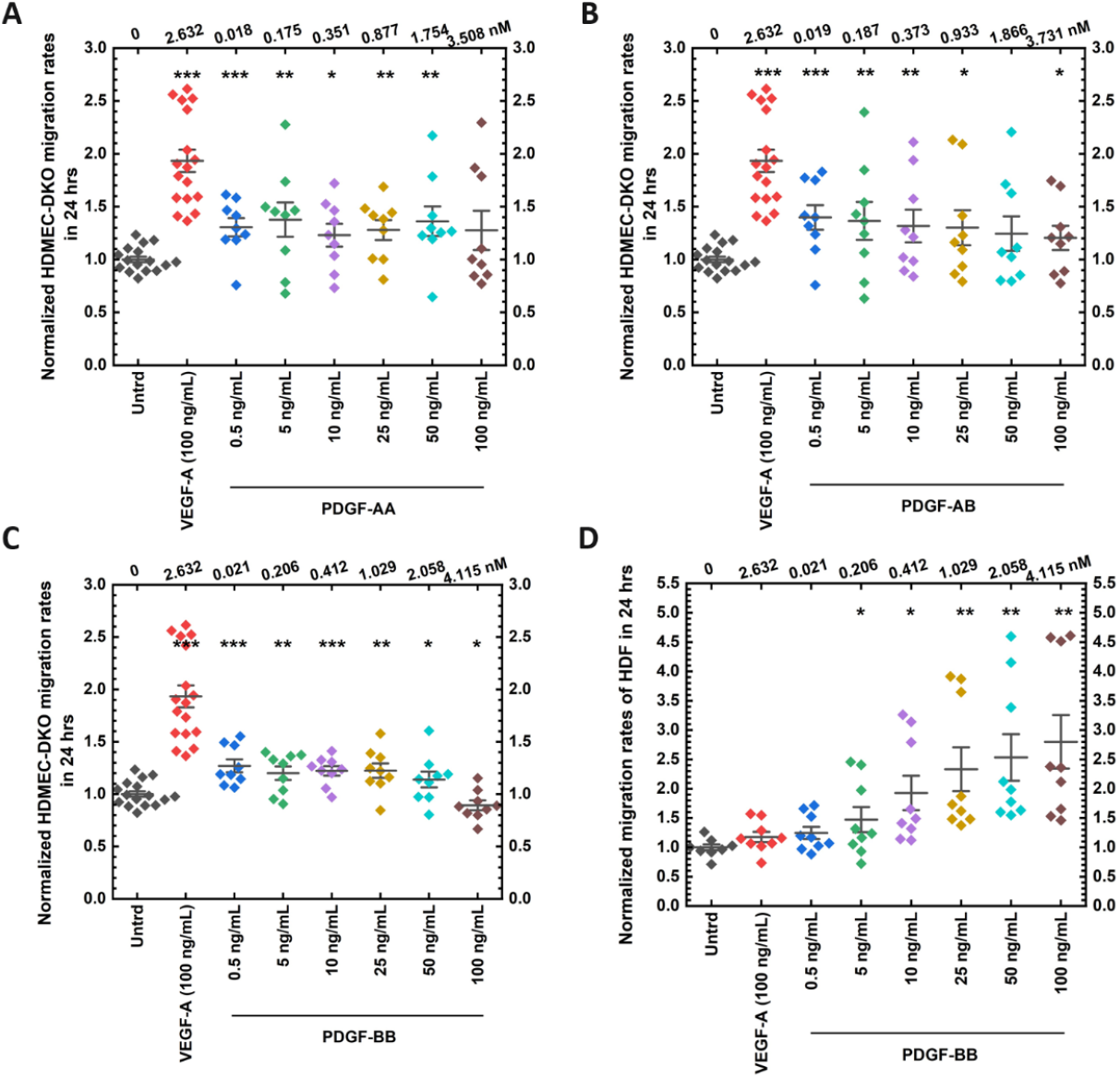
PDGFs stimulates migration in HDMECs (*PDGFRA*^*−/−*^ and *PDGFRB*^*−/−*^) and HDF. Migration rates after treatment of PDGFs were examined in (**A-C**) HDMECs (*PDGFRA*^*−/−*^ and *PDGFRB*^*−/−*^) and (**D**) in HDF. The concentrations of stimulating growth factors (ng/mL) are presented at the bottom of the plot, while the concentrations converted to the unit of pM are at the top of the plot. Three independent replicates were performed and analyzed. Error bars represent the standard error of mean (SEM). *p<0.05, **p<0.01, ***p<0.001 as determined by one-way ANOVA and multi-comparison post-hoc Tukey’s test.

### PDGF-induced EC responses are independent of VEGF-A upregulation

To determine whether treatment of PDGF-BB could have increased VEGF-A secretion by ECs and induced the PDGF-changes in angiogenic hallmarks [4,43], VEGF-A concentrations were measured in the cell culture supernatant following varying durations of PDGF-BB treatment. Our analysis revealed no statistically significant changes in VEGF-A levels between the untreated control and PDGF-BB-treated experimental cohorts (Figure 10B). Interestingly, within the VEGF-A positive subgroup, a progressive decline in VEGF-A concentration was noted over the course of incubation (Figure 10A). The trajectory of VEGF-A concentration reduction was accurately delineated by a regression curve (Figure 10C), with a calculated half-life of 32.33 minutes, suggesting the temporal stability of VEGF-A in the culture media after VEGF-A treatment.

**Figure 10.**
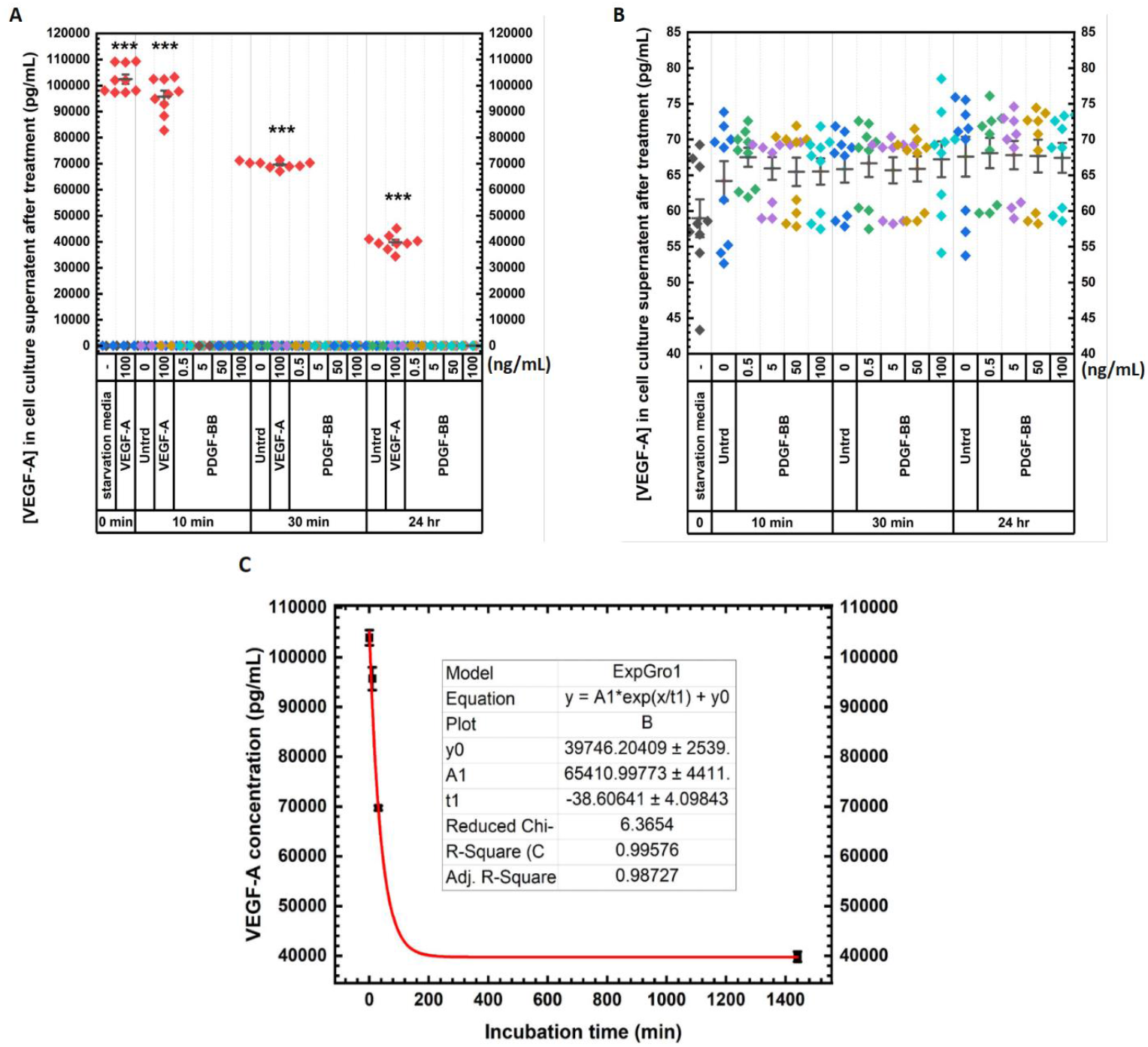
The concentration of VEGF-A exhibited no significant alteration upon PDGF-BB stimulation. **(A and B)** Subsequent to a 6-hour period of serum deprivation (1% FBS), HBMECs underwent treatment with growth factors as detailed. Post-incubation, levels of VEGF-A were assessed in the cell culture supernatant. **(C)** A fitting curve was generated to depict the correlation between VEGF-A concentration and in-vitro incubation duration. The determination of VEGF-A half-life time was computed utilizing the formula *t* = − *t*1 ⋅ ln 2, yielding a value of 26.76 minutes. Three independent replicates were performed and analyzed. Error bars represent the standard error of mean (SEM). *p<0.05, **p<0.01, ***p<0.001 as determined by one-way ANOVA and multi-comparison post-hoc Tukey’s test.

## Discussion

Our study elucidates the effects of cross-family PDGF-to-VEGFR binding in regulating endothelial signaling and functions. We found that (1) PDGFs played a critical role in augmenting EC angiogenic hallmarks, i.e., receptor and downstream protein phosphorylation, cell proliferation, and migration, through non-PDGF receptors, to a similar level compared to VEGF-A. (2) Low concentrations of PDGF-BB induced the angiogenic capacity of ECs to the greatest extent, whereas high concentrations showed a more potent effect in HDF. Other PDGFs did not exhibit such trends. (3) The angiogenic effects of PDGF-BB in HBMECs were more prominent than in HDMECs (*PDGFRA*^*−/−*^ and *PDGFRB*^*−/−*^). Other PDGFs showed similar impacts between EC types. (4) VEGF-A protein upregulation was not observed upon PDGF-BB stimulation, further validating the effects of PDGF-to-VEGFR signaling in ECs.

### PDGF-induced endothelial signaling and function

Cross-family interaction has been identified between PDGF and VEGFR as a novel angiogenic mechanism [4], the physiological functions are, however, yet unexplored. Here, we examined the angiogenic hallmarks of cross-family PDGF-to-VEGFR binding in regulating endothelial signaling and functions, namely receptor and effector activation, cell proliferation, and migration. We employed ECs that lack PDGF receptors (PDGFRs) on their plasma membranes to isolate the effects of possible PDGF-BB signaling through VEGFR2 from its signaling through PDGFR and also employed human fibroblasts that lack VEGFRs but have PDGFRs on their plasma membranes as a comparative control. To exclude the possible effects of upregulated VEGF-A transcription induced by PDGF treatment [31, 32], we also measured VEGF-A protein concentrations in the cell culture supernatant after varying durations of PDGF-BB treatment and confirmed that VEGF-A protein was not upregulated in response to PDGF-BB treatment.

### Low concentrations of PDGF showed more pronounced effects

Our study comprehensively examined a wide range of PDGF concentrations from 0.5 to 100 ng/mL and their effects on i) activation of angiogenic receptors VEGFR1 and VEGFR2, ii) endothelial cell proliferation, and iii) migration. We tested two concentrations of the PDGFs, 5 and 50ng/mL, for adaptor activation in the signaling pathway, as maximum stimulation of PDGF-BB was found at ≤50 ng/mL [33–35].

Our comprehensive analyses revealed that low concentrations of PDGFs induced EC functional changes at a higher level. Specifically, treatment of 0.5 ng/mL PDGF-AA or -BB induced the highest VEGFR1 phosphorylation concentration by approximately 2-fold in HBMECs, whereas high concentrations at 50**–** 100 ng/mL yielded the lowest receptor activation. PDGF-AA and -AB at 5 ng/mL activated downstream proteins, PLCγ1, Akt, and VEGFR2 at the investigated sites at a higher level than the concentration of 50 ng/mL. A low concentration of PDGF-BB at 0.5 ng/mL exhibited stronger activation of Akt and FAK compared to a high concentration at 50 ng/mL. Similar results were observed in the examinations of angiogenic hallmarks, where the highest impacts of PDGF treatment appeared at relatively low concentrations. Moreover, PDGF-BB showed half-maximal stimulation in HDFs at <10 ng/mL, which aligns with the literature [36, 37]. Notably, PDGF in human blood serum is measured to be 17.5 ng/mL [38], approximately 35-fold higher than 0.5 ng/mL, the concentration producing the most substantial angiogenic effects, which may explain why the functions of PDGF were long overlooked.

### PDGF-to-VEGFR1 signaling

Our observations indicate a strong connection between PDGF-induced signaling through VEGFRs, downstream proteins, and the angiogenic hallmarks of ECs. Following VEGFR2 activation by VEGF, downstream adaptors, PLCγ1, Akt, and FAK are activated [27] and play a role in regulating EC proliferation, survival, and migration [27–30]. In this study, we found that PDGFs could significantly activate VEGFRs and these proteins and induced EC angiogenic hallmarks. It is suggested that PDGF may signal through VEGFR1, VEGFR2, and downstream proteins to regulate EC angiogenesis.

Our observations suggest that ECs may engage in novel signaling pathways between PDGF and non-PDGF receptors on the plasma membrane. We propose that PDGFs induce VEGFR1 and VEGFR2 phosphorylation through VEGFR2 binding, since PDGF-to-VEGFR1 binding was not previously observed, but PDGF-to-VEGFR2 binding was seen [4]. In addition, VEGFR1 primarily heterodimerizes with VEGFR2 to regulate VEGF activity [2, 39]. Most VEGFR1 are in heterodimer forms with VEGFR2 on the plasma membrane of ECs [40]. These findings suggest the need to explore VEGFR1-VEGFR2 heterodimerization in PDGF-mediated cross-family interactions. Intriguingly, PDGF-AA and -BB were found to interact with NRP1 through phosphorylated VEGFR2, similar to VEGF-A_165_ interacting with NRP1 through VEGFR2[41, 42]. It suggests a possibility that NRP1 may also be involved in PDGF signaling on ECs through VEGFRs.

VEGFR1 has been typically considered a “decoy” receptor in angiogenesis due to its high binding affinity but limited kinase activity [43, 44]. However, an association between increased VEGFR1 levels and angiogenesis hallmarks was recently found [23, 26, 45–47], challenging the decoy paradigm. VEGFR1 protein expression is upregulated in tumors and ischemia, but VEGFR2 remains stable [23, 47]. VEGFR1 phosphorylation primarily activates PLCγ, PI3K, and MAPK, leading to cell survival and proliferation, whereas phosphorylation of VEGFR2 preferentially activates PLCγ, PKC, FAK, and PKB/Akt, regulating cell proliferation, migration, and permeability [27, 48]. Our studies are the first to demonstrate that PDGFs can activate VEGFR signaling in ECs, leading to significant phosphorylation of VEGFR1 and VEGFR2.

### PDGFs function variously in different cells

PDGF-BB showed more pronounced angiogenic effects, namely receptor activation, cell proliferation, and migration) in HBMECs compared with HDMECs (*PDGFRA*^*−/−*^ and *PDGFRB*^*−/−*^). Treatment with PDGF-BB led to a 220% increase in VEGFR1 phosphorylation in HBMECs, contrasting with a 180% increase in HDMECs (*PDGFRA*^*−/−*^ and *PDGFRB*^*−/−*^). Moreover, cell proliferation in HBMECs surged by 240% compared to 150% in HDMECs following PDGF-BB treatment, potentially influenced by the twofold higher concentration of VEGFR1 on the plasma membrane of HBMECs (700 ± 70 VEGFR1/cell) compared to HDMECs (*PDGFRA*^*−/−*^ and *PDGFRB*^*−/−*^) (300 ± 40 VEGFR1/cell) [21]. Moreover, heparin can inhibit growth factor effects on cell proliferation [49]. The medium culturing HDMECs (*PDGFRA*^*−/−*^ and *PDGFRB*^*−/−*^) contain a significant amount of heparin (0.0015 mg/mL).

The more remarkable effects of PDGF at higher concentrations in human fibroblasts were observed compared with the results in ECs with PDGF-BB at lower concentrations. PDGF-BB increased VEGFR1 phosphorylation by around 2-fold at 0.5 ng/mL in ECs and 8-fold at 100 ng/mL in human fibroblasts. Cell proliferation rate was elevated by 240% in HBMECs at 0.5 ng/mL compared with 280% in human fibroblasts by 100 ng/mL of PDGF-BB treatment. Cell migration was promoted to 130% enhancement by 0.5 ng/mL PDGF-BB in HBMECs and HDMECs (*PDGFRA*^*−/−*^ and *PDGFRB*^*−/−*^), whereas human fibroblasts reached the peak migration capacity by 100 ng/mL PDGF-BB of 2.8-fold increase. The data agreed with previous publications that strong cell proliferation induction only occurs with PDGF concentrations > 5 ng/mL and arises as the concentration increases [50]. PDGF-BB at 25 ng/mL was found to increase fibroblast proliferation by approximately 2-fold [51], comparable to our findings. The cell migration response is negligible at low concentrations and arises starting from 1 ng/mL [50]. PDGF-BB at 10 ng/mL increases fibroblast migration to around 200% [52], similar to our result. The less remarkable PDGF-induced impacts in ECs aligned with the differential receptor expression on these cells, as human fibroblasts express PDGFRs on the plasma membrane, but ECs do not. In such cases, PDGF displayed less activity on ECs than human fibroblasts due to the lack of plasma-membrane PDGFRs.

Our study highlighted significant VEGF-A-induced effects in HDFs lacking VEGFRs. VEGF-A significantly phosphorylated VEGFR1 and enhanced cell proliferation of HDFs, but not on cell migration. The validated impacts of VEGF-A in HDFs that lack VEGFRs but express PDGFRs on the plasma membrane, agreed with the published studies that VEGF-A could signal through PDGFRs [16] and demonstrated the effects of cross-family signaling between VEGF-A to PDGFRβ.

## Conclusion

By uncovering the angiogenic functions of PDGF-VEGFR interactions, we can better understand the cross-family angiogenic pathway, which may reveal alternate mechanisms for addressing vascular dysregulation. The pathways that downregulate VEGFR1 may be responsible for limiting excessive angiogenesis [53]. Proteomics is a reasonable attempt to systematically explore the underlying mechanisms. Our findings also provide essential data for computational modeling and systems biology to refine the signaling pathways.

## Acknowledgements

This material is based upon work supported by the National Science Foundation under Grant No. 2344705. Research reported in this publication was also supported by the National Institute of Heart, Lung, and Blood of the National Institutes of Health under Award Number 5R01HL159946-04. The funders had no role in study design, data collection and analysis, decision to publish, or preparation. Primary human dermal microvascular endothelial cells (HDMECs) (ATCC, CRL-4060) with PDGFRA and PDGFRB double knock out (DKO) via CRISPR/Cas9 were obtained at the Genome Engineering & Stem Cell Center at Washington University in St. Louis, MO.

## Author contributions

X.L., V.C., and P.I.I. conceived the experiments; X.L. conducted the experiments, analyzed the results, and prepared the figures and tables; S.K., S.G.N., and S.G. conducted the western blot examining VEGFR2 and adaptor phosphorylation in HDMEC-DKO and analyzed the results; Y.F. prepared Figure 3D–3G and Figure 4; X.L., Y.F., V.C., and P.I.I. contributed to the writing of the manuscript.

## Supplements

**Table S1.**
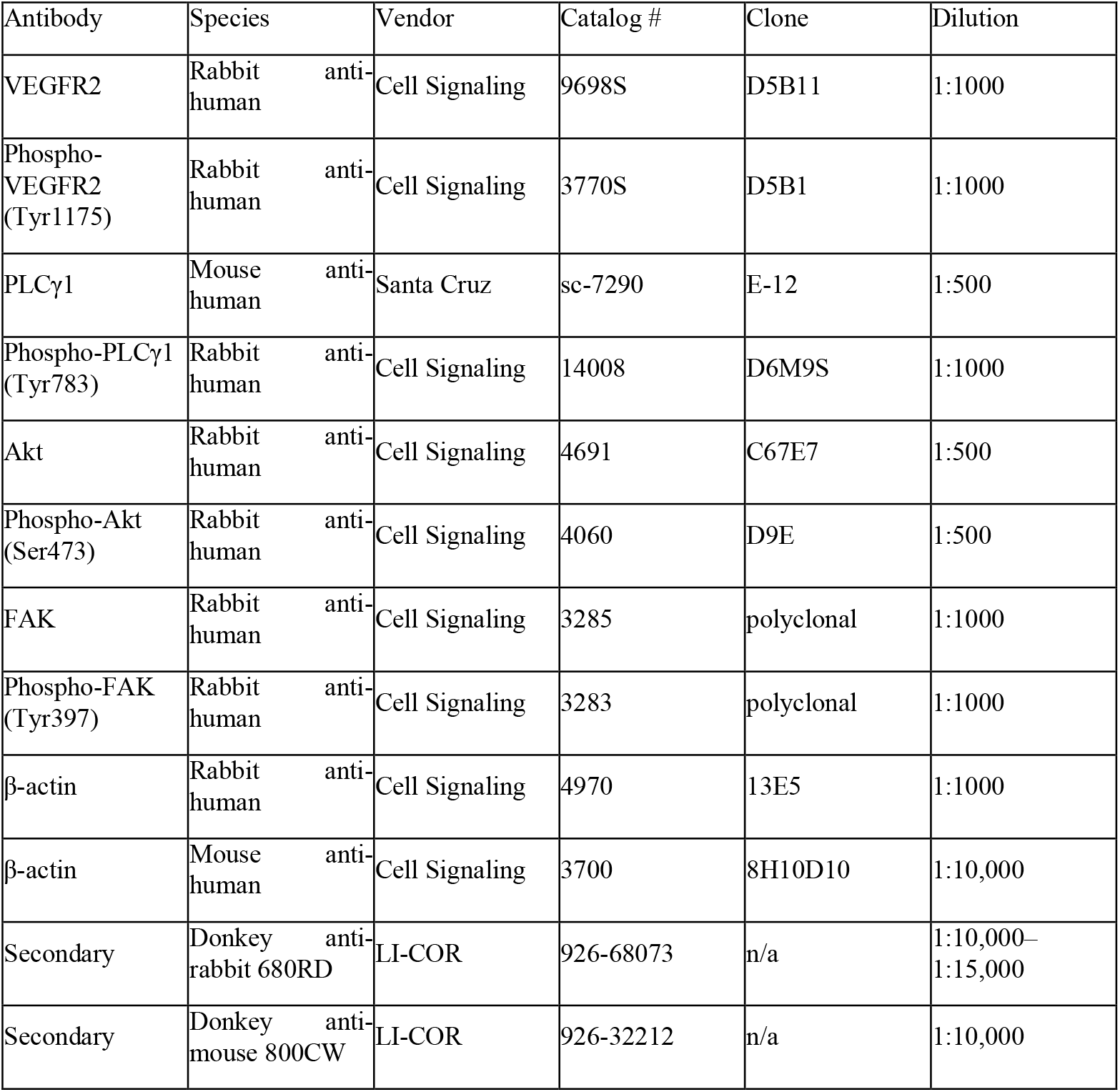
List of experimental antibodies.

