## Supplementary material for "Cross-family Signaling: PDGF Mediates VEGFR Activation and Endothelial Function": Suppl. Data

### Supplements

**Table S1. List of experimental antibodies**

| Antibody | Species | Vendor | Catalog # | Clone | Dilution |
|---|---|---|---|---|---|
| VEGFR2 | Rabbit anti-human | Cell Signaling | 9698S | D5B11 | 1:1000 |
| Phospho-VEGFR2 (Tyr1175) | Rabbit anti-human | Cell Signaling | 3770S | D5B1 | 1:1000 |
| PLCγ1 | Mouse anti-human | Santa Cruz | sc-7290 | E-12 | 1:500 |
| Phospho-PLCγ1 (Tyr783) | Rabbit anti-human | Cell Signaling | 14008 | D6M9S | 1:1000 |
| Akt | Rabbit anti-human | Cell Signaling | 4691 | C67E7 | 1:500 |
| Phospho-Akt (Ser473) | Rabbit anti-human | Cell Signaling | 4060 | D9E | 1:500 |
| FAK | Rabbit anti-human | Cell Signaling | 3285 | polyclonal | 1:1000 |
| Phospho-FAK (Tyr397) | Rabbit anti-human | Cell Signaling | 3283 | polyclonal | 1:1000 |
| β-actin | Rabbit anti-human | Cell Signaling | 4970 | 13E5 | 1:1000 |
| β-actin | Mouse anti-human | Cell Signaling | 3700 | 8H10D10 | 1:10,000 |
| Secondary | Donkey anti-rabbit 680RD | LI-COR | 926-68073 | n/a | 1:10,000–1:15,000 |
| Secondary | Donkey anti-mouse 800CW | LI-COR | 926-32212 | n/a | 1:10,000 |
